# MARRVEL-MCP: A Context-Engineered Natural-Language Query-to-Response Interface for Mendelian Disease Discovery

**DOI:** 10.1101/2025.11.26.690887

**Authors:** Zachary Everton, Jorge Botas, Seon Young Kim, Zhandong Liu, Hyun-Hwan Jeong

## Abstract

Rare disease variant interpretation requires navigating multiple genomic databases with strict input formats and synthesizing heterogeneous evidence, creating barriers for non-experts and cognitive burdens even for specialists. MARRVEL exemplifies this challenge by requiring precise queries (e.g., HGVS notation) and returning complex, difficult-to-synthesize outputs. To address this input-output asymmetry, we developed MARRVEL-MCP, a natural language interface enabling large language models (LLMs) to perform end-to-end variant interpretation via structured tool access.

This work demonstrates the impact of **context engineering**—the deliberate design of domain-aware tool environments and information scaffolding—in reshaping the role of model scale in genomics. MARRVEL-MCP equips LLMs with 39 tools spanning gene and variant utilities, pathogenicity databases, phenotype resources, expression atlases, ortholog data, and literature APIs. Without hard-coded pipelines, LLMs autonomously infer workflows, performing named-entity recognition, identifier normalization, and multi-database synthesis from narrative queries.

Using 45 expert-curated questions, lightweight models (3B–20B parameters) with MARRVEL-MCP matched or outperformed larger models lacking tool access. A 20B-parameter model (gpt-oss-20b) achieved a 95% pass rate versus 33% without MARRVEL-MCP, approaching state-of-the-art proprietary performance. Although accuracy remains below the 100% required for autonomous diagnosis and token usage is considerable, these results show that well-designed context can compensate for limited model capacity.

These findings establish context engineering as a core principle for biomedical AI, enabling efficient, scalable integration of LLMs with curated genomic resources. MARRVEL-MCP is available at https://github.com/hyunhwan-bcm/MARRVEL_MCP/.

## 1 Background

Determining the pathogenicity of genetic variants in rare Mendelian diseases requires integrating evidence across multiple dimensions: computational predictions of functional impact, population allele frequencies, clinical annotations from prior cases, gene expression patterns, model organism phenotypes, and supporting literature [1]. This multi-faceted assessment is essential because no single line of evidence is sufficient—a computationally predicted “damaging” variant found at high frequency in healthy populations is unlikely to be pathogenic, while a rare variant in a gene with matching model organism phenotypes and published case reports strengthens the clinical interpretation [2]. Yet this comprehensive evidence synthesis remains a bottleneck in rare disease diagnosis. Clinical geneticists must manually query multiple specialized databases, each with distinct input formats and data models, then cross-reference and reconcile outputs that may span different genomic coordinate systems, gene identifiers, and annotation standards. For research groups without dedicated bioinformatics support, this workflow can take hours per candidate variant and requires expertise in variant nomenclature standards, database provenance, and interpretation guidelines such as those from the American College of Medical Genetics and Genomics (ACMG) [1].

MARRVEL (Model organism Aggregated Resources for Rare Variant ExpLoration) [3] addresses part of this challenge by aggregating genomic, functional, and model-organism data into a unified platform. Since its release, MARRVEL has served more than 160,000 users and processed millions of queries. Furthermore, in 2025, MARRVEL recorded more than 43,000 active users worldwide in one year, becoming a widely adopted resource for rare disease research. By consolidating information from ClinVar [4], gnomAD [5], OMIM [6], expression databases, ortholog resources (DIOPT [7]), protein interaction networks (STRING [8]), and literature repositories, MARRVEL reduces the need to navigate 21 separate websites. However, MARRVEL itself operates within the traditional database paradigm: it expects users to provide precisely formatted inputs and returns comprehensive but heterogeneous outputs that still require significant manual interpretation.

This creates a fundamental **input-output asymmetry** that limits MARRVEL’s accessibility and efficiency. On the input side, users must convert their starting information—which might be an rsID from a sequencing report (e.g., rs193922679), a protein change from a clinical note (e.g., “p.Arg120Gln”), or a gene symbol with phenotype descriptor—into MARRVEL-compatible formats such as HGVS notation (e.g., NM_012160.3:c.541A>G) or genomic coordinates (e.g., chr6:99365567:T:C). This conversion requires knowledge of nomenclature standards, transcript selection, and coordinate systems (hg19 vs. hg38), creating a barrier to entry for clinicians and early-career researchers. On the output side, MARRVEL returns exhaustive annotations spanning pathogenicity predictions (dbNSFP [9]), clinical classifications (ClinVar [4]), population frequencies (gnomAD [5], DGV [10]), phenotype associations (OMIM [6], Geno2MP [11], HPO [12]), expression patterns (GTEx [13]), model organism data (DIOPT [7], FlyBase [14], WormBase [15], MGI [16]), and literature (PubMed [17])—all displayed on a single expansive page. Turning this into a clear answer to a clinical question (“Is this variant likely pathogenic in my patient with these symptoms?”) typically requires scrolling through sections, filtering irrelevant associations, cross-referencing allele frequencies against disease prevalence, verifying that model organism phenotypes align with the patient’s presentation, and conducting separate literature searches for mechanistic support. As a result, MARRVEL offers substantial workflow relief compared to querying each database independently, but it remains a tool optimized for expert users who can fluently navigate its structure and synthesize its outputs.

Large language models (LLMs) offer a complementary capability that could address this asymmetry. Unlike rule-based systems, LLMs can parse ambiguous natural language queries, extract relevant entities (gene symbols, phenotypes, variant descriptors), and generate human-readable summaries. [18] However, LLMs in isolation suffer from critical limitations for genomic applications: they cannot access current databases (their training data has a fixed cutoff), they may hallucinate gene-phenotype associations or variant frequencies, and they cannot guarantee their reasoning follows valid biological workflows. A model asked “Is variant X pathogenic?” might generate a plausible-sounding answer (“Variant X is classified as likely pathogenic based on computational predictions…”) without actually checking ClinVar or any other authoritative source. Tool-use frameworks address this by enabling LLMs to invoke external functions. The Model Context Protocol (MCP) [19] provides a standardized interface for exposing database APIs, computational tools, and retrieval systems to LLMs through structured function calls. When a user asks a question, the LLM can issue a sequence of tool invocations—such as converting an identifier, querying a database, and retrieving literature—rather than generating an answer from parametric memory alone. This grounds the output in real, current data while preserving the LLM’s ability to synthesize information into natural language responses.

However, simply providing tool access is insufficient for reliable genomic analysis. Recent systems have explored LLM-database integration in bioinformatics, including retrieval-augmented generation over biomedical literature [20, 21, 22], code generation for analysis scripts [23], and conversational interfaces to single databases [24]. These efforts demonstrate that LLMs can interact with biological data sources, but they have not addressed the core challenge of **multi-database workflow orchestration** in the absence of hard-coded pipelines. For instance, determining variant pathogenicity requires not just querying ClinVar, but first resolving identifiers, then querying multiple annotation sources (e.g., dbNSFP and gnomAD), cross-referencing with gene-phenotype databases (e.g., OMIM and HPO), and finally synthesizing evidence according to domain constraints (e.g., a high gnomAD frequency contradicts pathogenicity regardless of computational predictions). Executing this correctly requires more than tool availability—it requires **context engineering**.

We define context engineering as the deliberate construction of structured, domain-aware information environments that shape how models interpret queries and compose operations. In contrast to prompt engineering, which provides behavioral guidance through natural language instructions, context engineering operates at the level of the reasoning environment itself: the tools available, their semantic descriptions, the format of their inputs and outputs, intermediate state representations, and the relationships between data elements. For variant interpretation, effective context includes not just database access, but also structured knowledge of variant nomenclature standards (HGVS, rsIDs, protein notation), genomic coordinate systems and liftover procedures (hg19 ↔ hg38), ontology-aware phenotype relationships (HPO term hierarchies), and provenance metadata that enables the LLM to reason about evidence quality (ClinVar’s assertion criteria, gnomAD’s population stratification, dbNSFP’s score distributions). This contextual scaffolding allows the LLM to remain adaptive—generating novel reasoning traces for diverse queries—while operating within scientifically grounded constraints that prevent common failure modes such as mixing coordinate systems, querying databases with malformed identifiers, or synthesizing evidence without respecting ACMG interpretation logic.

Context engineering differs from standard retrieval-augmented generation (RAG), which retrieves relevant text passages to augment prompts, but does not provide executable tools or enforce structured workflows. It also differs from general-purpose agent frameworks like AutoGPT [25] or ReAct [26], which provide planning scaffolds but lack the domain-specific tools and constraints necessary for biological reasoning. A general agent might generate a plan like “Step 1: Search for variant pathogenicity. Step 2: Summarize findings”—but without access to ClinVar’s API and knowledge of HGVS notation, it cannot execute this plan reliably. Context engineering instead embeds domain knowledge directly into the tool environment, enabling specialized but flexible reasoning within a defined operational scope.

To demonstrate this principle, we developed **MARRVEL-MCP**, an MCP-based tool environment that enables LLMs to execute end-to-end variant interpretation from natural-language requests. Given a question like “What is known about variant rs193922679 and its disease relevance?”, an LLM equipped with MARRVEL-MCP autonomously assembles and executes the required workflow: converting the rsID to genomic coordinates, querying dbNSFP for functional predictions, retrieving ClinVar annotations, checking gnomAD allele frequencies, and synthesizing the evidence into a structured summary—all without requiring the user to specify the analysis steps or intermediate formats. Importantly, MARRVEL-MCP does not rely on hard-coded pipelines or explicit workflow specifications. When provided only with tool descriptions such as `convert_rsid_to_variant: “Converts an rsID to chromosome, position, reference and alternate alleles”` and `get_variant_dbnsfp: “Retrieves functional prediction scores from dbNSFP given genomic coordinates”`, the LLM autonomously infers the correct sequencing through semantic reasoning: it recognizes that pathogenicity assessment requires genomic coordinates, observes that the user provided an rsID, identifies the conversion tool, invokes it, and uses the returned coordinates for subsequent queries. This emergent workflow construction—where valid multi-step analyses are composed from atomic operations without explicit chaining rules—demonstrates that carefully designed context can enable adaptive reasoning while maintaining operational reliability.

In this architecture, context engineering becomes a first-class design principle. The availability of specific tools (39 in total, spanning identifier conversion, pathogenicity databases, phenotype resources, expression atlases, ortholog comparisons, and literature APIs), the structure of their intermediate outputs (standardized JSON with explicit field semantics), provenance-aware evidence (each result tagged with database name and version), and ontology support (HPO term expansion, OMIM phenotype mappings) collectively guide the LLM’s reasoning trajectory. This controlled contextual scaffolding allows the model to remain adaptive and transparent—users can inspect the tool call sequence—while operating within scientifically grounded constraints. The result is a system that balances the expressive flexibility of LLMs with the rigor and trustworthiness required for biomedical decision support.

Our contributions are threefold, collectively demonstrating that **context engineering is as important as model scale for practical genomic AI systems**:

### 1. Natural-language to genomic workflow execution

We develop an MCP tool suite that allows an LLM to translate free-form user intent into automatically executed variant-interpretation pipelines, removing both the pre-query formatting burden and post-query manual triage. The system handles diverse entry points—rsIDs, HGVS notation, protein changes, gene symbols with phenotypes—and autonomously constructs valid analysis workflows without hard-coded decision trees.

### 2. Context engineering can outweigh model scale

We show that 3B–20B parameter models equipped with MARRVEL-MCP achieve accuracy comparable to or exceeding 50B+ parameter models operating without tool access. On a benchmark of 45 expert-curated questions, a 20B-parameter model with MARRVEL-MCP reached 95% accuracy compared to 33% without tools—demonstrating that well-designed tool environments and domain context can compensate for limited model capacity in specialized tasks. This has direct implications for cost-efficiency: smaller models with structured context are both more capable and orders of magnitude cheaper to deploy than frontier-scale models.

### 3. A benchmark for future genomic tool-using agents

We curate and release 45 realistic benchmark tasks covering rare-disease gene discovery, variant interpretation, phenotype association queries, and literature synthesis scenarios. This provides a standardized evaluation resource for future MCP-based genomic assistants and enables direct comparison of different context engineering approaches.

Together, these results establish a template for connecting LLM agents to curated biomedical databases: rather than relying solely on ever-larger general-purpose models, it is often more effective and cost-efficient to connect modest-sized LLMs to well-engineered, domain-specific tool environments. In the context of rare disease research, MARRVEL-MCP illustrates how this pattern can transform a mature but complex database into an interactive, intent-driven assistant that lowers barriers to sophisticated genomic analysis while preserving MARRVEL’s curated evidence as the authoritative backbone and maintaining the essential role of expert oversight.

## 2 Implementation

MARRVEL-MCP connects the flexibility of large language models (LLMs) to the reliability of curated biomedical databases through the Model Context Protocol (MCP), enabling structured and iterative access to trusted genomic resources (Figure 1). Through this protocol, LLMs send requests to a tool endpoint hosted on an MCP server instance, which retrieves the requested information via MARRVEL or other integrated databases such as PubMed and the Human Phenotype Ontology. The MCP server processes each query by interpreting its semantic intent, issuing corresponding database requests, and filtering and condensing the resulting data to minimize token usage before returning optimized, structured context to the model.

**Figure 1.**
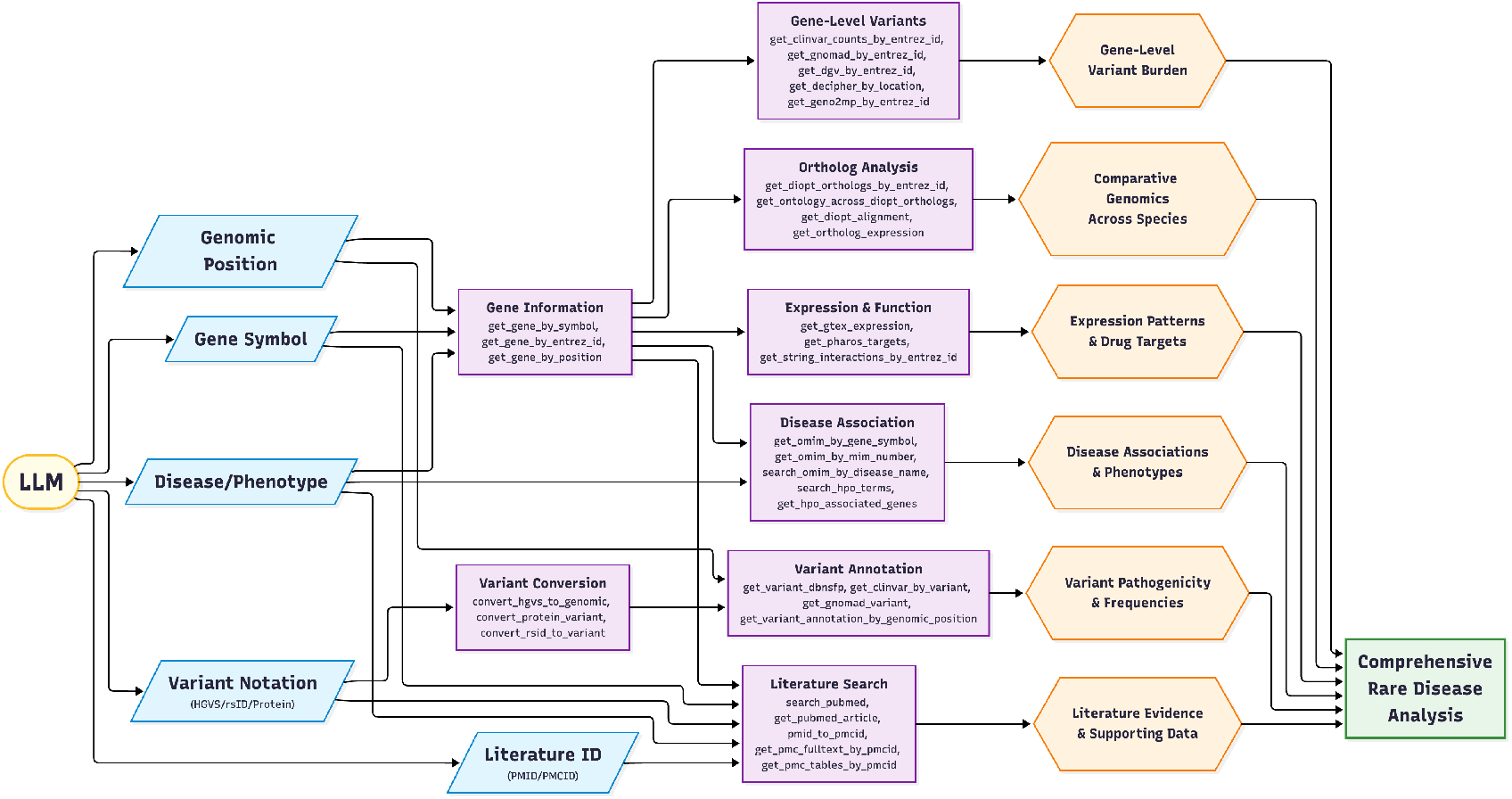
Overview of the MARRVEL-MCP tool-chain and downstream functionality, illustrating how an LLM orchestrates multiple entry points (genomic position, gene symbol, disease/phenotype, variant notation, and literature ID) to activate interconnected analytical modules. These modules include gene-level variant analysis, ortholog comparison, expression and functional profiling, disease association mapping, variant annotation, and literature mining. The integrated outputs converge into a unified pipeline enabling comprehensive rare disease analysis, supporting detailed interpretation of variant pathogenicity, cross-species genomics, phenotypic associations, and evidence-based insights.

**Figure 2.**
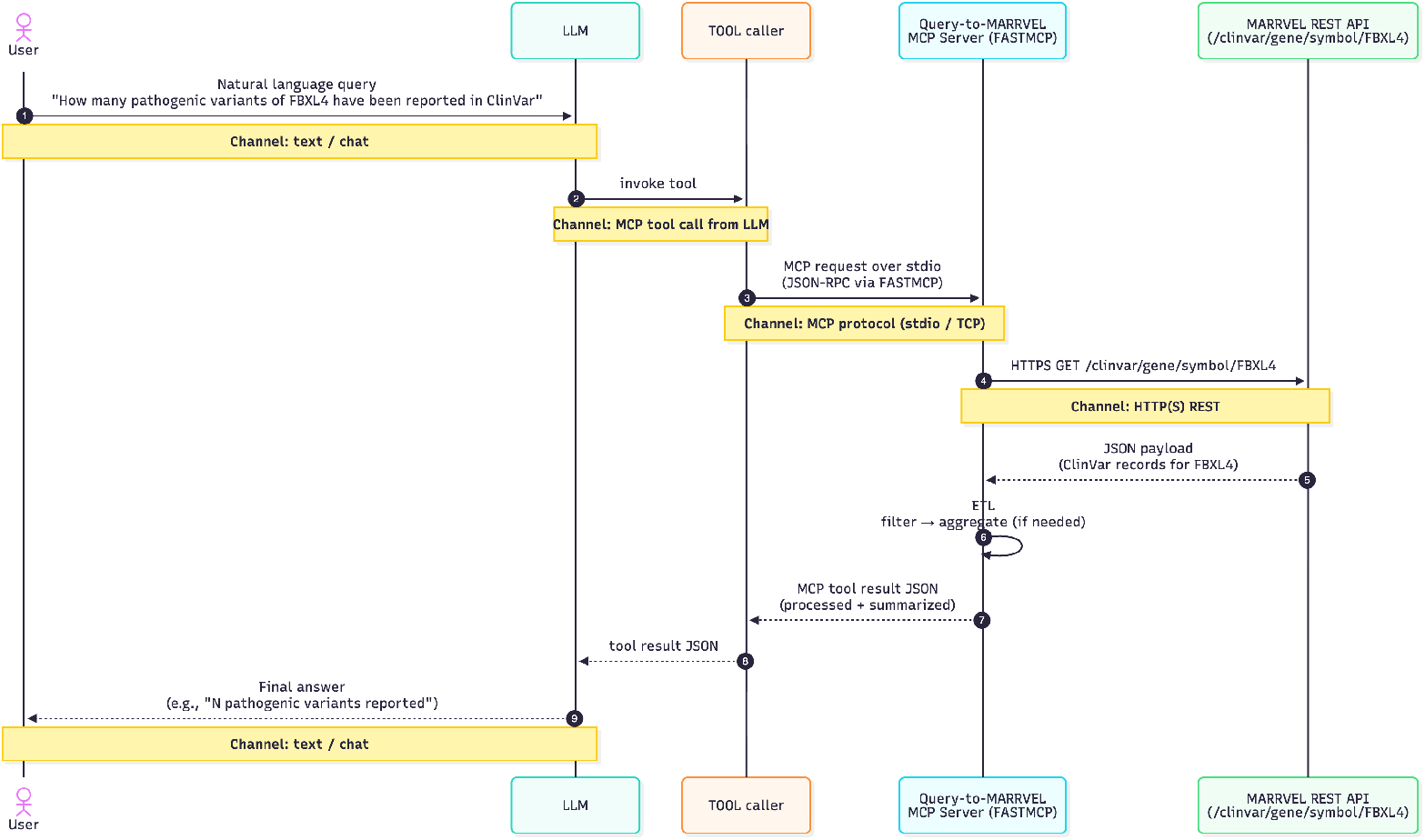
A sequence diagram illustrating how a natural-language query from a user (e.g., “How many pathogenic variants of FBXL4 have been reported in ClinVar?”) flows from the LLM to an MCP tool caller, which sends a JSON-RPC request to a FASTMCP server. The server queries the MARRVEL REST API, performs optional ETL/aggregation on the returned ClinVar data, and returns a summarized JSON result back through the tool caller to the LLM, which delivers the final text answer to the user.

As the LLM generates responses, it may iteratively issue multiple tool calls, using intermediate results to formulate subsequent queries, thereby enabling dynamic resolution of complex, multi-layered biological questions that would traditionally require multiple independent searches. This orchestration allows diverse entry points — including genomic position, gene symbol, disease or phenotype, variant notation, and literature identifiers — to trigger interconnected analytical modules such as gene-level variant analysis, ortholog comparison, expression and functional profiling, disease association mapping, variant annotation, and literature mining (Figure S1). The integrated outputs are consolidated into a unified analytical pipeline that supports comprehensive rare disease interpretation, including detailed assessment of variant pathogenicity, cross-species genomic context, phenotype-genotype associations, and evidence-based literature support.

To address the variety of gene identifiers and information sources that are commonly encountered when describing genes and variants, we developed thirty-nine MCP tools enabling large-language models to interface with the MARRVEL database and other web APIs via REST and GraphQL endpoints. These tools were implemented using the Python version of Fast-MCP framework, which provided a streamlined and standardized mechanism for exposing our query functionality to LLM workflows (Figure S1). As a result, we created 39 different MCP tools for LLMs that interface with MARRVEL and other web APIs (Table 1).

**Table 1:** Overview of available tool categories and their functionalities in MARRVEL-MCP.

| Category | Description | Number of Tools |
| --- | --- | --- |
| Gene Utility | Gene symbols (across naming conventions) and genome position | 4 |
| Variant Utility | Matchups between variant notations | 4 |
| Liftover | Perform liftovers between hg19/GRC37 and hg38/GRC38 | 2 |
| dbNSFP | Variant impact predictions, scores and rank scores from different sources | 1 |
| ClinVar | Provides clinical information on recorded variants (benign, pathogenic, etc) | 2 |
| gnomAD | Provides counts and statistics on variants found in the gnomAD control database | 2 |
| DGV | Provides counts and statistics on variants found in the DGV control database | 1 |
| DECIPHER | Search DECIPHER for control variants and variant statistics in a genomic region | 1 |
| geno2MP | Links rare genotypes to human phenotype ontology terms | 1 |
| OMIM | Search OMIM for gene-phenotype information by gene, variant, mim number or disease name | 4 |
| HPO | Search Human Phenotype Ontology by phenotype or HPO term | 2 |
| STRING | Search STRING DB for interactions between genes | 2 |
| DIOPT | Search across model organisms for ortholog genes, Gene Ontology Terms, domain info and sequence alignment | 3 |
| Expression | Retrieves tissue and cell type-specific expression levels for a given gene across human and other model organisms, as well as drug targeting information from Pharos | 3 |
| PubMed | Perform searches and return components of PubMed/PubMed Central publications by PMID or PMCID | 11 |

### 2.1 Gene and Variant utility tools

One of the largest limitations of MARRVEL is its limited support for different gene and variant symbol formats. To remedy this, MARRVEL MCP includes eight tools designed to convert the gene/variant in the format provided to the format required for other lookups.

For genes, we include tools to look up gene information by genome location, HUGO Gene Nomenclature Committee (HGNC) symbol, or Entrez gene ID. Each of these query types will return entries with these four fields, allowing the LLM to convert between one of the formats to any of the other three.

For variants, we include tools to extract chromosome, position, reference sequence and alternate sequence from different variant notations. The `convert_hgvs_to_variant` tool extract these fields from Human Genome Variant Society (HGVS) formats, including cDNA, genomic and protein notation. The `convert_rsid_to_variant` tool works similarly, returning the chr/pos/ref/alt info for a given NCBI dbSNP [27] rsID number. The last two tools work in opposite directions: `convert_protein_variant` extracts chr/pos/ref/alt info from a given gene and amino acid change, and `get_variant_annotation_by_genomic_position` returns the amino acid change given the allele and position.

### 2.2 Liftover tools

When examining genomic sites or regions, it is also important for the AI model to be able to perform liftovers between hg19/GRCh37 and hg38/GRCh38. The two liftover tools included provide this functionality, and can be used by the LLM when necessary.

While developing the MCP tools, we observed that the LLM will often have difficulty determining when to convert between hg19 and hg38 coordinates when working with a given tool’s output. Because of this, each tool is configured to take and return hg38 coordinates, even if the original database the tool queries uses hg19. Liftover tool calls are used within other tools if the coordinates need to be converted to hg19 to query an external database or the external database returns hg19 coordinates.

### 2.3 Variant prioritization tools

Many questions regarding genetic variants involve the variant’s impact. As such, we have 7 tools focused on assessing variant impact with various databases. dbNSFP is one database referenced, a comprehensive resource containing many functional variant predictions. The `get_dbNSFP_by_entrez_id`tool returns scores, predictions and rank scores for many different models, including AlphaMissense [28], CADD [29], MCAP [30], among others. The tool also calculates the average rank score for all models, which can be useful if the user is less familiar with the individual models referenced.

While dbNSFP contains predicted variant effect, ClinVar is a database that contains actual variant effect. This database has two different methods - one method returns the pathogenicity grade for a given variant, and the other tool returns information at the gene level. To reduce the number of tokens given to the LLM, the gene level tool only returns entry counts for each classification, returning the number of variants marked “benign”, “likely benign”, “likely pathogenic”, and “pathogenic”. More information is provided by the variant level tool.

DECIPHER is another database of pathogenic variants, focusing on variants found in individuals with rare genetic diseases. For DECIPHER, we implemented one tool that queries pathogenic information given a genome location. This tool is similar to ClinVar’s variant tool, but because of this database’s structure, we can search by a genomic region rather than a single base pair. Another difference is that ClinVar is generalized, while DECIPHER is focused on rare diseases.

Another two databases focus on healthy control populations instead of pathogenic cases. GnomAD has two tools implemented (one queries at the gene level and the other at the variant level) and DGV has one variant-level tool. Both of these databases are useful in testing to see if a given variant is being selected against.

### 2.4 Gene-phenotype database tools

Other may focus on phenotype associations at the gene level instead of the variant level. Adding onto the tools listed above, we implemented tools for querying Geno2MP database, and four tools for querying the OMIM database. The Geno2MP database is populated directly from genomic centers and can be useful to discover new gene-phenotype links. To access this database, MARVELL MCP includes a tool that counts the frequency of different types of mutations recorded in a given gene.

OMIM is another gene-phenotype database, focusing more on associations supported by published literature. We included two tools that investigate gene-phenotype associations from the genetics end, with one tool that will return OMIM entries and associated phenotypes for a gievn gene and one tool that will return entries and phenotypes for a given variant. Because OMIM focuses more on phenotypic information than Geno2MP, we are also able to query from the other end: one tool will perform a general search for OMIM pheotype entries given a discription of the phenotype, and one tool will pull up the specific OMIM entry for a given phenotype using the phenotype’s mim number (an accession ID used by OMIM). In this way, researchers can look for phenotypes connected to their gene of interest of genes connected to their phenotype of interest depending on where their project has started.

The Human Phenotype Onotology (HPO) database is the third gene-phenotype database, this one focused more on phenotypes. DECIPHER and OMIM are among the databases used to create the database, so there is not as much it brings on the genetic side, but it is much more organized on the phenotype side and thus is useful from that angle. The `search_hpo_terms` tool allows the LLM to search HPO ids and descriptions for their phenotype of interest, and the `get_hpo_associated_genes` term allows the LLM to retrieve genes linked to a given phenotype. Between the three databases, the model has multiple options to check for relationships between genes, variants and phenotypes.

### 2.5 Drug Targeting

When a pathogenic gene or variant is discovered, researchers often want to know how to target it. For this purpose, MARRVEL MCP includes one tool that looks up drug interactions on the Pharos database [31]. Given a gene, the tool will query the database and return a list of known ligands and their effectiveness.

### 2.6 Expression database tools

An important part of human genetics research is identifying where the gene is typically expressed, a question answered by gene expression databases. GTEx is a widely used expression database that provides gene expression levels for many types of human tissue. The get_gtex_expression tool allows LLMs to consult GTEx, returning the median expression level in each recorded tissue type for a given gene.

The gene expression atlas can be used to take expression analysis a step further, examining expression across different stages of development and across gene orthologs in other species. The `get_ortholog_expression` MCP tool lets LLMs query the number of expression annotations in each tissue/cell function/development stage for a given gene’s orthologs in a specified organism. In this way, the LLM can check the expression of genes between different organisms.

### 2.7 Gene orthologs

Gene orthologs themselves are also an area of interest for genetics researchers. By accessing information on DIOPT, LLMs can find orthologous genes across all model organisms, as well as conserved protein domains and sequence alignments. The first tool, `get_orthologs_by_entrez_id`, returns a list of ortholog gene ids along with the taxon ids of the organisms these genes are found in. The second tool, `get_ontology_by_entrez_id`, instead returns Gene Ontology terms associated with the gene’s orthologs in a given organism. The last tool, get_diopt_alignment, shows the sequence alignment for all orthologs of a given gene id, as well as UniProt information on conserved domains.

### 2.8 Gene association database tools

When multiple genes are found to be connected to a certain phenotype, a common validation step is to check if the genes are related to each other through pathways, regulation, or other ways. STRING, a database of protein associations, can be used to query associations between genes and is made accessible LLMs through two MARRVEL MCP tools. The first tool, `get_string_interactions_by_entrez_id`, lists all genes associated with the given gene along with the association type and strength. The second tool, `get_string_interactions_between_entrez_ids`, returns any STRING associations between the two genes along with the association types and strengths. By using these tools, the LLM model can look for other genes involved with a gene of interest, or see if two genes of interest are connected in some way.

### 2.9 Literature tools

Researchers commonly rely on PubMed, which indexes article citations and abstracts, and PubMed Central, which provides access to full-text publications, to gather background information on genes of interest. Within MARRVEL-MCP, LLMs can leverage the `search_pubmed` MCP tool to retrieve PubMed IDs (PMIDs) relevant to a natural-language query. Given a PMID, the system can obtain higher-level article metadata via `get_pubmed_article` or convert it to a corresponding PubMed Central ID (PMCID) using `pmid_to_pmcid`. Once a PMCID is available, the model can selectively access different article components—including abstracts, full text, tables, or figure captions.

This modular retrieval framework not only supports precise, context-aware evidence gathering but also establishes the foundation for the next generation of MARRVEL-MCP, which will expand article-level reasoning, integrate richer biomedical ontologies, and unify literature insights with structured database outputs.

### 2.10 Software Tools, Library, and benchmark reproducibility

In this work, we employed the following software tools and libraries:

- Python [32] for core scripting, data processing, and experiment.
- FastMCP [33] as our Model Context Protocol (MCP) framework to expose Python tools to language models in a structured, high-performance way.
- GraphQL [34] for schema-based, strongly typed data access, enabling flexible and efficient querying of our back-end services.
- REST [35] as the underlying architectural style for stateless HTTP APIs, supporting interoperability across services.
- LangChain [36] as the main orchestration framework for building LLM-powered workflows and tool-augmented agents.
- Publiplots [37] to generate publication-quality, Python-based visualizations with a clean, modular plotting API.
- OpenRouter [38] as a unified API layer to access and route between multiple large language models.
- llama.cpp [39] for efficient local inference of large language models on commodity hardware.

All benchmark results and the code used to generate the figures are publicly available at: https://github.com/LiuzLab/MARRVEL_MCP_manuscript

## 3 Results

### 3.1 MARRVEL-MCP enables natural language variant interpretation through autonomous workflow construction

To assess whether MARRVEL-MCP can bridge the gap between natural-language questions and structured genomic analysis, we tested its ability to handle two representative query types: a simple identifier-based lookup and a complex narrative case description. These experiments demonstrate that MARRVEL-MCP not only executes database queries but autonomously constructs valid multi-step workflows from minimal user input.

#### 3.1.1 Structured variant annotation from a single natural language query

We queried Claude Haiku 4.5 integrated with MARRVEL-MCP using the instruction: “summarize all available variant functional prediction rank scores for rs193922679 in a table format.” This example reflects a common real-world scenario encountered by clinical geneticists, where a single rsID derived from a sequencing report serves as the initial reference point for evaluating predicted functional impact, with results expected to be presented in a structured format suitable for clinical interpretation. Notably, this is a technically challenging request, as accurately interpreting such a query requires a thorough understanding of specialized genomics terminology and domain-specific jargon, which may not be immediately intuitive without appropriate background knowledge.

As shown in Figure 3A, the MARRVEL-MCP agent successfully decomposed this request into the required computational workflow. The agent first recognized that functional prediction scores from dbNSFP require genomic coordinates rather than rsIDs. It autonomously invoked `convert_rsid_to_variant` to translate rs193922679 into X-154031253 T>A in MECP2 (hg38) Figure S3, then called `get_variant_dbnsfp` with these coordinates to retrieve pathogenicity predictions from AlphaMissense, CADD, REVEL, and other models. The agent returned a concise table reporting both rank scores and qualitative assessments (e.g., “Deleterious (D)” for MutationTaster and CADD rank score 0.99), formatted exactly as a geneticist would need for variant interpretation. From the user’s perspective, the entire workflow was triggered by a single sentence, without any need to specify coordinate systems, database endpoints, or output formatting.

**Figure 3.**
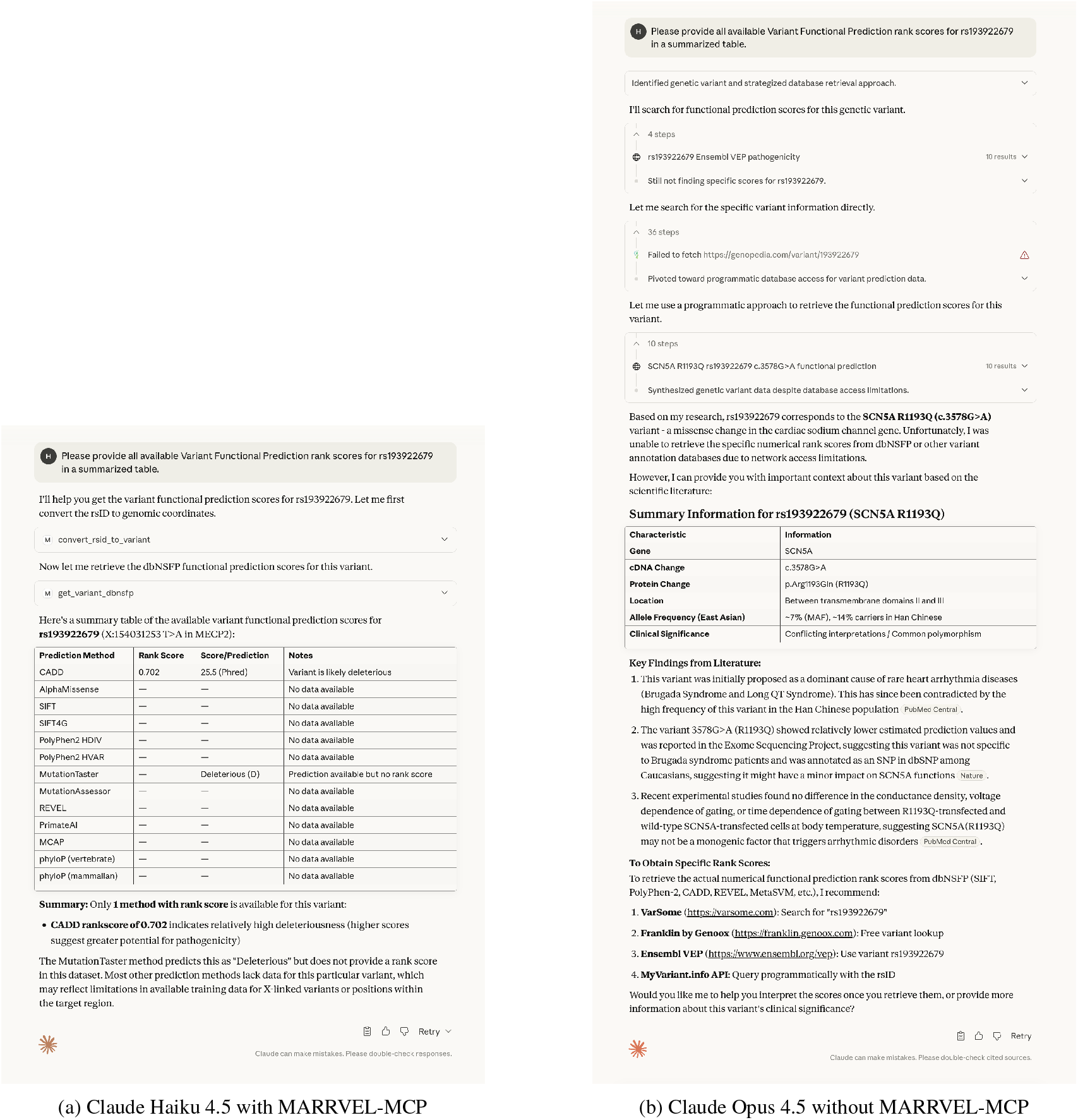
Structured variant annotation from natural language: MARRVEL-MCP enables precise answers while general-purpose models provide narrative summaries. **(A)** Claude Haiku 4.5 equipped with MARRVEL-MCP successfully interpreted the query “all available Variant Functional Prediction rank scores for rs193922679 in a summarized table,” autonomously converting the rsID to genomic coordinates (X-154031253 T>A) via `convert_rsid_to_variant`, then querying dbNSFP functional predictions via `get_variant_dbnsfp`. The system returned a structured table with rank scores and qualitative assessments (e.g., CADD rank score 0.72, “Deleterious” for MutationTaster) formatted for clinical review. Importantly, this workflow emerged from tool descriptions alone, without hard-coded pipelines—the model inferred that dbNSFP requires coordinates and selected the appropriate conversion tool. **(B)** In contrast, Claude Opus 4.5 tested without MARRVEL-MCP but with all standard capabilities enabled (web search, extended reasoning) produced a qualitatively different response. Opus incorrectly identified rs193922679 as SCN1A R1193Q, retrieved biological context via a web search, but failed to return the requested structured table of computational predictions. This comparison demonstrates that even state-of-the-art general-purpose models struggle to produce analytically complete, database-grounded outputs without access to domain-specific tools, highlighting the importance of context engineering for practical genomic applications.

Importantly, this workflow was not hard-coded. When we examined the tool-call trace, we observed that the model received only tool descriptions such as `convert_rsid_to_variant: “Converts an rsID to chromosome, position, reference and alternate alleles”` and `get_variant_dbnsfp: “Retrieves functional prediction scores from dbNSFP given genomic coordinates (hg38)”`. The correct sequencing—rsID conversion must precede coordinate-based queries—emerged from semantic reasoning about tool input requirements rather than from explicit workflow rules. This demonstrates that carefully designed context (tool availability, input/output specifications, coordinate system constraints) enables LLMs to construct valid analysis pipelines autonomously.

We contrasted this result with Claude Opus 4.5, currently the most capable model in the Claude family, tested with all standard capabilities enabled including web search and extended reasoning (Figure 3B). Despite its substantially larger capacity, Opus without MARRVEL-MCP access produced a qualitatively different response. The model incorrectly searched the web for information about rs193922679 and identified that the variant is associated with SCNS1A R1193Q, while Haiku correctly identified the region on MECP2 (Figure S3). However, it returned a narrative summary of web search results rather than the structured table of computational predictions that the user requested. Opus retrieved the incorrect biological context and could not access the specific dbNSFP functional scores or format them appropriately. This highlights that even state-of-the-art general-purpose models struggle to produce analytically complete, structured responses without access to domain-specific tools—reinforcing the importance of context-aware systems that ground LLM reasoning in authoritative databases.

This comparison demonstrates two key principles. **First, context engineering enables emergent capabilities**. By providing tool descriptions with explicit input/output semantics (e.g., “requires hg38 coordinates”) and constraint information (e.g., dbNSFP scores are coordinate-based), MARRVEL-MCP allowed a lightweight model to autonomously infer the correct workflow without hard-coded decision trees. **Second, structured tool access can outweigh raw model capacity for domain-specific tasks**. Haiku (roughly one-fifth the inference cost of Opus) delivered the more useful answer in this genomic context, demonstrating that well-designed tool environments can make smaller models competitive with—or superior to—much larger general-purpose systems when evaluated on task-specific utility rather than general intelligence.

#### 3.1.2 Automated workflow construction from narrative clinical text

To assess MARRVEL-MCP’s performance on unstructured clinical narratives, we used a case description adapted from a published study that employed MARRVEL in its findings [40]: *“For the gene HEXB, assess whether the variants c*.*298delC and G473S, found in a compound-heterozygous state, are likely to be pathogenic and sufficient to explain a Sandhoff-disease-like presentation. Provide a concise conclusion*.*”*

This query differs fundamentally from the rsID example in that it embeds multiple biomedical entities within narrative text, uses mixed variant notations (HGVS cDNA and protein nomenclature), and asks for a synthesized interpretation rather than raw data retrieval. Answering correctly requires extracting structured entities, normalizing identifiers, querying multiple databases, and integrating evidence according to domain logic.

As shown in Figure 4, MARRVEL-MCP autonomously performed these steps using Gemini-2.5-Flash. The system extracted the gene symbol (HEXB), two sequence variants (c.298delC, G473S), the inheritance pattern (compound heterozygosity), and the phenotype descriptor (Sandhoff-disease-like presentation). It then constructed and executed a workflow spanning six tools: `get_gene_by_symbol` to retrieve the Entrez ID for HEXB, `hgvs_to_genomic` to convert c.298delC from cDNA notation to genomic coordinates, `convert_protein_variant` to resolve G473S to a genomic position, `clinvar_by_variant` and `gnomad_by_variant` to assess pathogenicity and population frequency for each variant, and `search_pubmed` to retrieve supporting literature on HEXB and Sandhoff disease. The final response integrated these data sources into a conclusion addressing the user’s question about disease causation.

**Figure 4.**
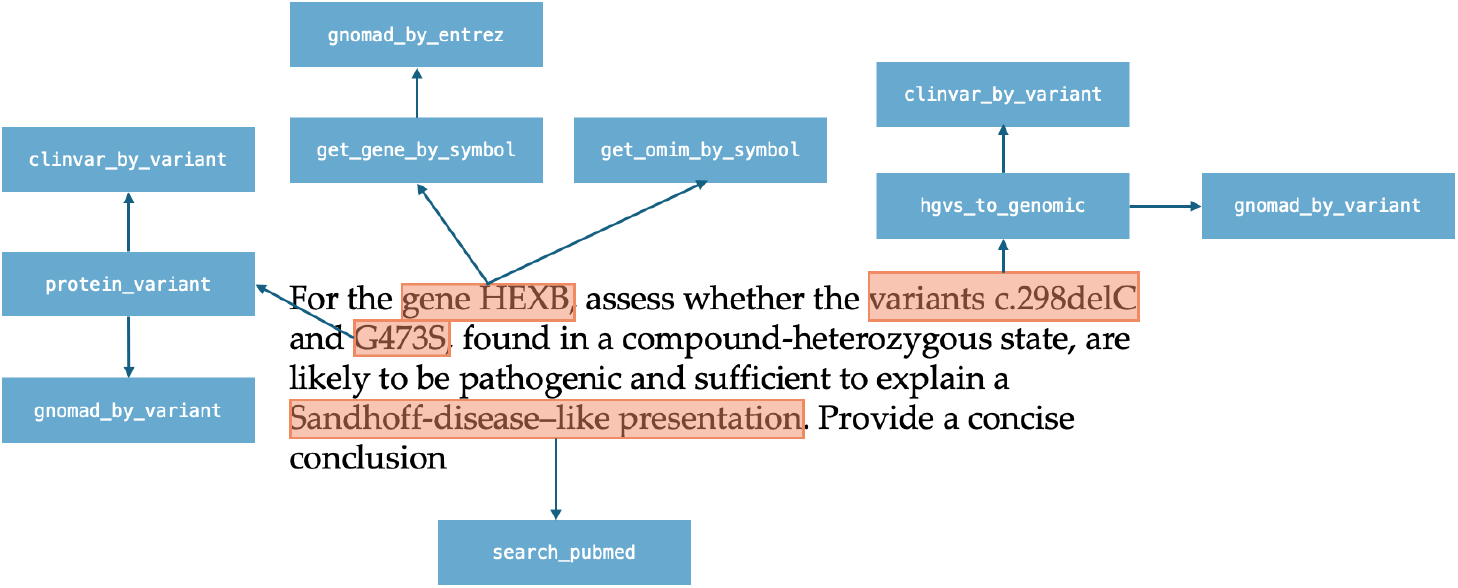
Automated workflow construction from narrative clinical text. MARRVEL-MCP autonomously parsed a complex clinical case description—*”For the gene HEXB, assess whether the variants c*.*298delC and G473S, found in a compound-heterozygous state, are likely to be pathogenic and sufficient to explain a Sandhoff-disease-like presentation”*—and constructed a six-step analysis workflow without manual entity annotation or workflow specification. The system performed named-entity recognition to extract the gene symbol (HEXB), two variants in different notations (HGVS cDNA: c.298delC; protein: G473S), inheritance pattern (compound heterozygosity), and phenotype (Sandhoff disease). It then autonomously selected and sequenced appropriate tools: `get_gene_by_symbol` to retrieve the Entrez ID, `hgvs_to_genomic` and `convert_protein_variant` to normalize variant identifiers, `clinvar_by_variant` and `gnomad_by_variant` to assess pathogenicity and population frequency, and `search_pubmed` for supporting literature. This example demonstrates that MARRVEL-MCP extends beyond simple database queries by integrating unsupervised biomedical NER, semantic identifier normalization, and goal-directed evidence synthesis—enabling direct operationalization of narrative variant interpretation statements from clinical genetics practice.

This example demonstrates three capabilities that distinguish MARRVEL-MCP from simple database query interfaces. **First, unsupervised named-entity recognition:** the system identified structured biomedical elements within free text without requiring the user to tag entities or specify their types. **Second, semantic normalization:** the system recognized that c.298delC requires HGVS-to-coordinate conversion while G473S requires protein-to-coordinate conversion, and selected appropriate tools for each. **Third, goal-directed synthesis:** rather than returning raw database outputs, the system recognized that the query’s implicit goal (“are [these variants] likely to be pathogenic and sufficient to explain [this phenotype]”) required integrating ClinVar annotations, population frequencies, and phenotype associations into a coherent assessment.

Together, these examples show that MARRVEL-MCP can operate directly on the language of clinical genetics—turning narrative case descriptions into executable analyses without manual encoding of entities or workflows—while preserving MARRVEL’s curated evidence as the authoritative data source. (Figure S4)

### 3.2 Small models with MARRVEL-MCP match or exceed large models without tools

Equipping lightweight models with MARRVEL-MCP substantially improved their accuracy on variant interpretation tasks, enabling smaller models to match or exceed the baseline performance of much larger systems. We evaluated this effect across nine LLMs ranging from 3B to 235B parameters using a benchmark of 45 expert-curated questions answerable using MARRVEL data. Each model was tested in two conditions: without MARRVEL-MCP access (baseline) and with full access to all 39 tools. Responses were evaluated by Claude Sonnet 4 as an external judge, with pass rate defined as the proportion of questions judged correct.

MARRVEL-MCP improved accuracy for every model tested, with particularly pronounced gains for smaller systems (Figure 5). The ministral-3b model increased from 18% baseline accuracy to 44% with tools—a 2.4× improvement. The effect was even more striking for mid-sized models: qwen3-14b improved from 22% to 80%, while gpt-oss-20b achieved 95% accuracy with MARRVEL-MCP compared to just 33% without—a nearly 3× gain. Critically, gpt-oss-20b with MARRVEL-MCP reached performance comparable to claude-sonnet-4 (100%), the strongest model in our benchmark, despite having an estimated 10× fewer parameters. This demonstrates that structured access to domain tools can compensate for limited model capacity: a 20B-parameter model with MARRVEL-MCP substantially outperformed a 120B-parameter model (gpt-oss-120b: 33% baseline) operating without tool access.

**Figure 5.**
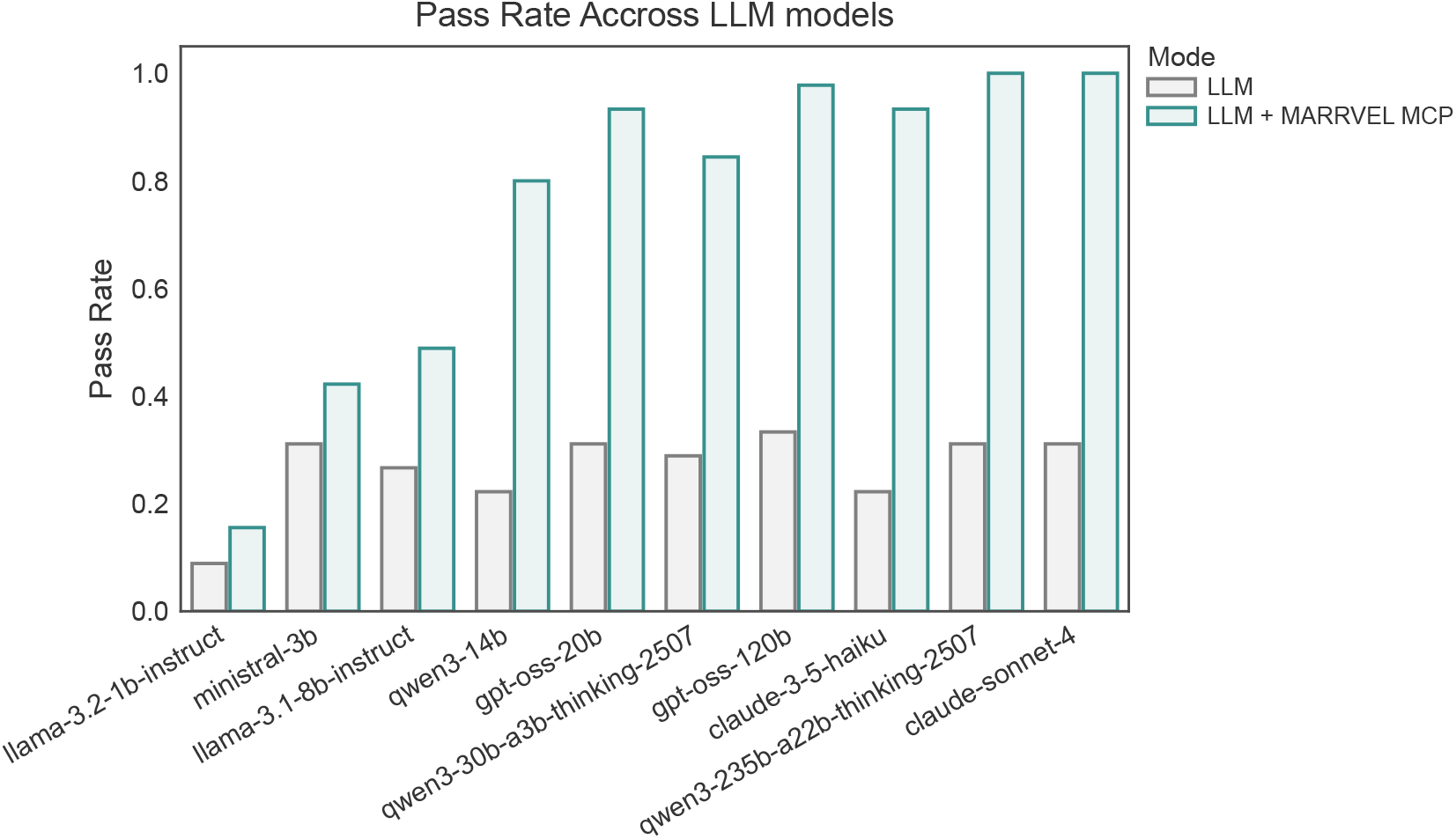
MARRVEL-MCP improves accuracy across all model sizes, with especially pronounced gains for lightweight models. We evaluated nine LLMs ranging from 3B to 235B parameters on 45 expert-curated questions answerable using MARRVEL data. Each model was tested without MARRVEL-MCP access (light bars) and with full access to all 39 tools (dark bars). Pass rate was defined as the proportion of answers judged correct by Claude Sonnet 4. Enabling MARRVEL-MCP substantially improved accuracy for every model, with gains ranging from 2.4× (ministral-3b: 18% → 44%) to nearly 3× (gpt-oss-20b: 33% → 95%). Critically, several small-to-mid-sized models with MARRVEL-MCP (qwen3-14b: 80%, gpt-oss-20b: 95%) matched or exceeded the baseline performance of much larger models without tools (gpt-oss-120b: 33%). Error bars represent 95% confidence intervals computed via Wilson score intervals for binomial proportions. This demonstrates that structured access to domain tools via context engineering can compensate for limited model capacity, enabling smaller models to achieve accuracy comparable to frontier-scale systems on domain-specific tasks.

The relationship between model size and accuracy reveals that tool access fundamentally reshapes the performance landscape (Figure 6). Without MARRVEL-MCP, accuracy scales roughly with model size: baseline performance ranges from 18% (llama-3.2-1b) to 35% (gpt-oss-120b), following an approximately log-linear trend. With MARRVEL-MCP enabled, models cluster near the upper accuracy range regardless of size. Even 3B–14B models achieved 44–80% accuracy with tools, approaching the 95–100% ceiling occupied by the strongest systems. The smallest model with MARRVEL-MCP (ministral-3b: 44%) outperformed the largest model without it (gpt-oss-120b: 33%). This suggests that for genomics tasks grounded in structured databases, context engineering and tool orchestration can be more important determinants of practical utility than raw model scale.

**Figure 6.**
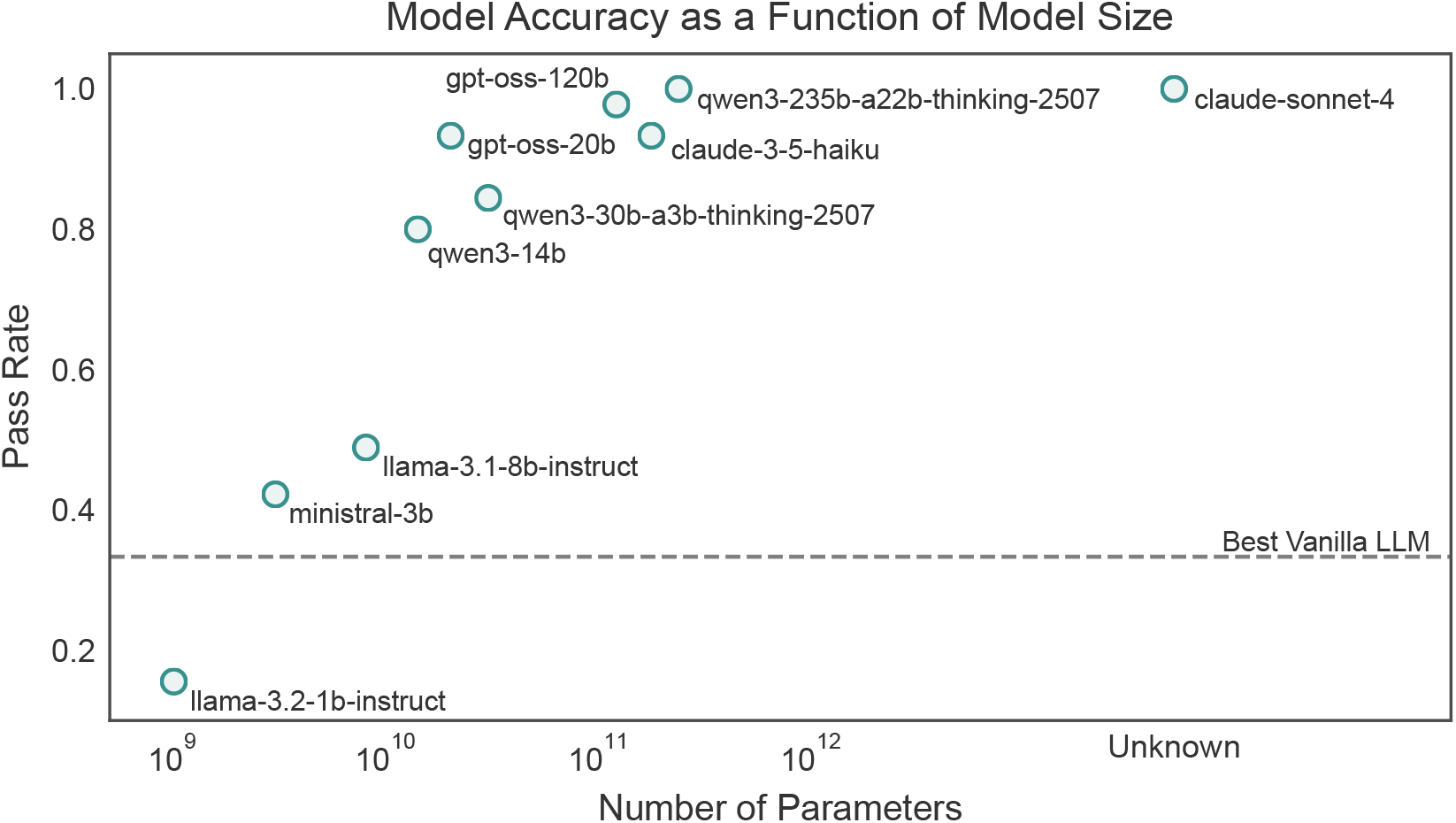
Tool access compresses the relationship between model size and accuracy, enabling small models to approach frontier performance. Pass rates for models equipped with MARRVEL-MCP are plotted against reported parameter counts. Without tool access (not shown), accuracy scales roughly log-linearly with model size, ranging from 18% (3B parameters) to 35% (120B parameters). With MARRVEL-MCP enabled, models cluster near the upper accuracy range (44–100%) regardless of size. The smallest model with tools (ministral-3b: 44%) outperformed the largest model without tools (gpt-oss-120b: 33%, indicated by dashed reference line). Notably, gpt-oss-20b with MARRVEL-MCP (95%) reached performance comparable to claude-sonnet-4 (100%), despite an estimated 10× difference in parameter count. Parameter sizes for proprietary models (Claude series, GPT-OSS) are not publicly disclosed and are indicated as “Unknown.” This pattern suggests that for genomics tasks grounded in structured databases, context engineering and tool orchestration can be more important determinants of practical performance than raw model scale, with direct implications for cost-efficient deployment.

However, this accuracy gain comes with a token cost that scales inversely with model capability (Figure 7). Smaller models required more elaborate reasoning chains to reach correct answers, consuming substantially more tokens than larger models performing the same tasks. For example, ministral-3b averaged approximately 15,000 tokens per question with MARRVEL-MCP enabled, compared to roughly 2,000 tokens for claude-sonnet-4—a 7.5× difference. This pattern held across the model range: llama-3.1-8b consumed median 12,000 tokens per question, while qwen3-30b required only 3,000. The token overhead arises from two sources. First, verbose tool specifications: each question triggers the transmission of all 39 tool descriptions (averaging 8,000 tokens) to establish context. Second, longer reasoning traces: smaller models often require multiple attempts, backtracking, or redundant tool calls to reach correct answers, whereas larger models navigate the tool space more efficiently.

**Figure 7.**
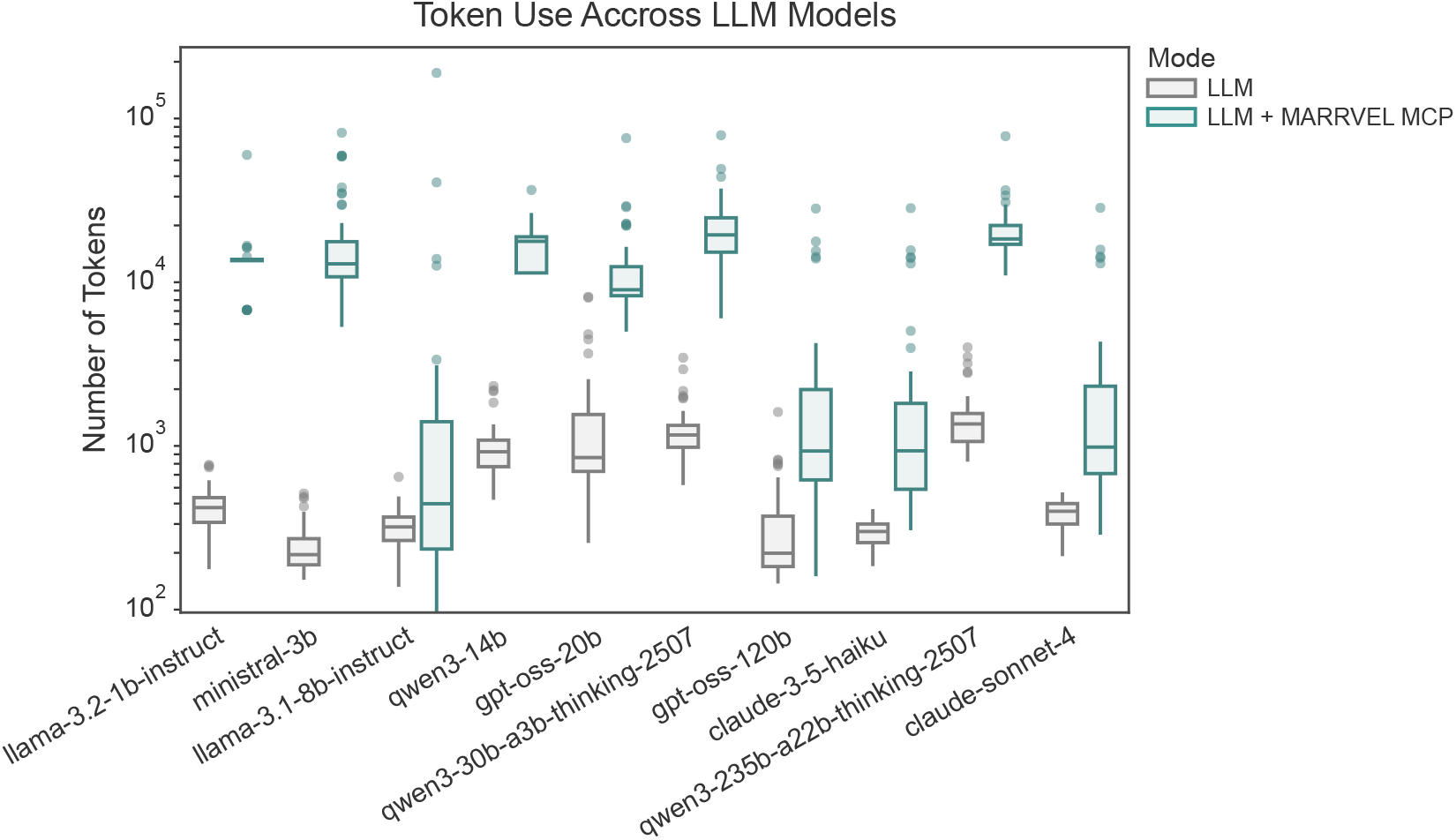
Token usage scales inversely with model capability, creating a cost-accuracy tradeoff. We measured total token consumption (input + output) per question for each model with and without MARRVEL-MCP access. Boxplots show median, quartiles, and outliers across the 45-question benchmark. Smaller models required substantially more tokens than larger models when using MARRVEL-MCP: ministral-3b consumed median 15,000 tokens per question compared to 2,000 for claude-sonnet-4—a 7.5× difference. This overhead arises from two sources: (1) verbose tool specifications transmitted with each query (approximately 8,000 tokens for 39 tool descriptions), and (2) longer reasoning chains in which smaller models require multiple attempts, backtracking, or redundant tool calls to reach correct answers. Without MARRVEL-MCP (light boxes), all models consume relatively few tokens (200–500) because they generate answers from parametric memory without invoking external tools. This pattern has direct implications for deployment: while MARRVEL-MCP makes small models more accurate (Figure 5), the computational savings from using a 3B versus 30B model are partially offset by 4–5× higher per-query token costs. At current API pricing, mid-sized models (14B–30B parameters) offer the best accuracy-cost tradeoff for production use, achieving 80–95% accuracy with manageable token overhead (3,000–5,000 tokens per query).

This cost structure has direct implications for deployment. While MARRVEL-MCP makes smaller models substantially more accurate, the computational savings from using a 3B model versus a 30B model are partially offset by 4–5× higher token consumption per query. At current API pricing, a 3B model with MARRVEL-MCP may cost more per query than a 30B model with MARRVEL-MCP, despite lower per-token rates. For high-throughput applications (e.g., batch processing of clinical cohorts), this suggests that mid-sized models (14B–30B) offer the best accuracy-cost tradeoff. For latency-sensitive or low-volume use cases (e.g., interactive clinical support), the absolute token cost remains acceptable even for small models.

### 3.3 Error analysis reveals addressable limitations

To understand the failure modes of MARRVEL-MCP and identify opportunities for improvement, we manually reviewed all incorrect responses from gpt-oss-20b, the best-performing open model (95% accuracy, 2/45 failures). We categorized errors into three types based on their underlying cause.

#### Tool selection errors (1/2 failures)

The model invoked an inappropriate tool for the query context. In one case, when asked “What HPO terms are associated with developmental delay?”, the model called `get_gene_by_symbol` with “developmental delay” as input rather than recognizing this as a phenotype descriptor requiring `search_hpo_terms`. This suggests that tool descriptions, while sufficient for most queries, may benefit from explicit negative examples (e.g., “Do not use this tool for phenotype queries—use search_hpo_terms instead”) or few-shot demonstrations of correct tool selection for ambiguous entity types.

#### Incomplete evidence synthesis (1/2 failures)

The model retrieved correct data but failed to integrate it according to ACMG interpretation logic. When asked to assess pathogenicity for a variant, the model correctly queried ClinVar (classification: “Uncertain Significance”) and gnomAD (allele frequency: 0.0001), but concluded “likely pathogenic” without acknowledging the conflicting ClinVar annotation or applying ACMG criteria for reconciling discordant evidence. This failure mode suggests that current tool outputs, while structured, do not include sufficient provenance metadata or interpretation guidelines. Augmenting tool responses with mini-tutorials (e.g., “ClinVar ’Uncertain Significance’ should not be upgraded to ’Likely Pathogenic’ without strong functional evidence—see ACMG guidelines”) could improve synthesis quality.

We also examined pass rate by query category across all models. The 45 questions span four types: variant pathogenicity assessment (n=15), gene-phenotype associations (n=12), cross-species ortholog queries (n=10), and literature synthesis (n=8). Claude Sonnet 4 with MARRVEL-MCP achieved 100% accuracy on variant pathogenicity and ortholog queries but 87.5% (7/8) on literature synthesis tasks. Manual inspection revealed that the failed literature query required integrating information across three PubMed abstracts to answer a comparative question (“How do loss-of-function phenotypes in flies compare to mouse models for this gene?”), whereas the model summarized each abstract independently without cross-referencing. This suggests that unstructured text integration remains more challenging than database retrieval even with tool access, and points to a need for explicit multi-document reasoning scaffolds or retrieval strategies optimized for comparative synthesis.

These error patterns are addressable through targeted interventions. Tool selection errors can be reduced via supervised fine-tuning on MARRVEL-MCP interaction logs, allowing models to internalize common query → tool mappings. Evidence synthesis failures suggest augmenting tool outputs with domain constraints (e.g., ACMG rules, ontology hierarchies) that guide interpretation. Literature integration challenges may benefit from retrieval-augmented generation techniques that explicitly structure multi-document comparisons. Importantly, none of these failure modes reflect fundamental limitations of the tool-use paradigm—they represent opportunities for refinement through context compression, fine-tuning, or enhanced provenance metadata.

### 3.4 Clinical context: MARRVEL-MCP as decision support, not autonomous diagnosis

While 95% accuracy represents substantial improvement over baseline LLMs (33% for gpt-oss-20b), it remains below the threshold required for autonomous clinical use. Variant interpretation in clinical genetics demands near-perfect accuracy because misclassification can lead to incorrect treatment decisions, inappropriate genetic counseling, or failure to identify causative variants in diagnostic odysseys that may span years [1]. The American College of Medical Genetics and Genomics (ACMG) guidelines for variant interpretation emphasize that evidence integration requires expert judgment, particularly when computational predictions, population frequencies, and functional data provide conflicting signals [1]. A 5% error rate, while acceptable for exploratory research or candidate prioritization, is incompatible with autonomous diagnostic workflows where mistakes carry clinical consequences.

Current performance positions MARRVEL-MCP as a **decision-support tool** that accelerates expert workflows rather than replacing expert judgment. In this role, MARRVEL-MCP offers clear value: it eliminates the manual formatting, database navigation, and evidence retrieval steps that consume hours of a geneticist’s time per candidate variant, allowing experts to focus on the synthesis and interpretation tasks where human judgment is most critical. For example, a clinical geneticist can ask “Summarize pathogenicity evidence for BRCA1 c.5266dupC in a patient with early-onset breast cancer,” receive a structured summary of ClinVar annotations, gnomAD frequencies, functional predictions, and relevant literature within seconds, and then apply their expertise to contextualize this evidence within the patient’s phenotype and family history. This workflow is substantially more efficient than querying each database independently, while preserving the expert’s authoritative role in final interpretation.

This positioning aligns with broader trends in clinical AI, where decision-support systems that augment rather than replace expert judgment have demonstrated the most sustainable impact [41]. Future work should focus on improving accuracy to 98–99% through supervised fine-tuning on expert-annotated interaction logs, which would make MARRVEL-MCP suitable for higher-stakes applications such as automated evidence packaging for clinical review. Even at current performance levels, however, the system provides meaningful value by transforming hours of manual database work into seconds of natural-language interaction, while maintaining the expert geneticist as the final arbiter of clinical interpretation.

## 4 Discussion and Conclusion

MARRVEL-MCP demonstrates that structured tool environments can reshape the relationship between model scale and genomic task performance. A 20B-parameter model with MARRVEL-MCP achieved 95% accuracy compared to 33% without tools—matching systems 5–10× larger and approaching the performance of state-of-the-art proprietary models. This pattern held consistently: lightweight models with appropriate context outperformed much larger systems operating without tool access. The practical implication is direct: modest-sized models with well-designed context can deliver reliable genomic analysis at a fraction of the cost of frontier alternatives, challenging the assumption that biomedical AI progress depends primarily on ever-larger models.

This effect stems from context engineering—encoding domain constraints, nomenclature standards, coordinate systems, and provenance metadata directly into the tool environment. By providing explicit tool input requirements and structured intermediate outputs, MARRVEL-MCP enables even small models to autonomously construct valid multistep workflows without hard-coded pipelines. The rs193922679 example crystallizes this principle: Haiku with MARRVEL-MCP delivered structured dbNSFP predictions formatted for clinical review by autonomously inferring the correct workflow (rsID conversion, coordinate-based query, tabular summary), while a much larger model without tools provided only narrative web search results. Task utility depends not solely on model intelligence but on whether the reasoning environment makes correct execution possible and incorrect execution detectable.

The 95% accuracy achieved by our best-performing open model, while substantially better than baseline, remains insufficient for autonomous clinical deployment. Variant interpretation demands near-perfect accuracy because errors have direct patient consequences. The ACMG guidelines emphasize that evidence integration requires expert judgment, especially when computational predictions conflict with population data or clinical annotations. Our error analysis revealed instructive failure modes: incorrect tool selection for ambiguous entity types and incomplete evidence synthesis where models failed to apply ACMG reconciliation logic when integrating conflicting annotations. These failures point to addressable improvements—tool descriptions augmented with negative examples, outputs enriched with domain constraints, and explicit encoding of interpretation guidelines—but underscore that expert oversight remains essential.

The appropriate role for MARRVEL-MCP is decision support that accelerates expert workflows rather than replacing judgment. A geneticist can pose a natural-language query and receive a structured evidence summary within seconds, then contextualize this within patient phenotype and family history. The workflow becomes: ask, review, interpret—eliminating hours of manual database navigation and identifier formatting while preserving the expert’s authoritative role. This positioning aligns with successful clinical AI systems, which demonstrate sustainable impact through augmentation rather than autonomy. Future work should focus on improving accuracy to 98–99% through supervised fine-tuning on expert-annotated interaction logs, making MARRVEL-MCP suitable for higher-stakes applications like automated evidence packaging for clinical review.

MARRVEL-MCP’s accuracy gains come at a computational cost that reveals fundamental design tradeoffs. Smaller models required 4–7× more tokens than larger models to reach correct answers, consuming median 15,000 tokens per query versus 2,000 for the most capable systems. This overhead arises from transmitting all tool descriptions with each query and from longer reasoning traces involving backtracking and redundant tool calls. At current API pricing, this creates a context efficiency paradox: a 3B model with MARRVEL-MCP may cost more per query than a 30B model with MARRVEL-MCP despite lower per-token rates. The current implementation conflates what the model needs to know—tool semantics and domain logic—with what must be retransmitted each time—tool schemas and specifications. Several architectural solutions could address this: tool indexing to load only relevant subsets, macro-tools encapsulating common workflows, supervised fine-tuning to internalize tool semantics, and context caching for static schemas. The optimal architecture likely combines these approaches, and identifying the right balance between generality, efficiency, and maintainability remains an important research question.

Our 45-question benchmark covers representative variant interpretation tasks but does not span the full complexity of clinical genetics: complex inheritance patterns, structural variants, somatic mutations, or pharmacogenomics. Future benchmarks should expand to these domains and include adversarial cases with incomplete or conflicting database information. MARRVEL-MCP provides database access but does not explicitly encode ACMG interpretation logic; models must infer that high population frequency contradicts pathogenicity or that annotations require contextualization. Integrating structured guidelines—either through augmented tool outputs or explicit interpretation modules—would improve synthesis quality. While MARRVEL-MCP includes literature tools, our benchmark emphasized database queries, and the single comparative literature question revealed that multi-document synthesis remains more challenging than structured retrieval. Explicit scaffolding for comparative analysis across abstracts would address this limitation. Finally, clinical deployment would require addressing regulatory considerations (FDA clearance as Software as a Medical Device), integration with EHR and laboratory systems, and institutional review processes—translational challenges beyond our current technical scope.

We have released MARRVEL-MCP as an open resource at https://github.com/hyunhwan-bcm/MARRVEL_MCP/, providing a reference implementation for integrating LLM agents with curated biomedical databases. The design patterns demonstrated here—explicit constraint encoding, provenance-aware outputs, ontology integration, and emergent workflow construction—generalize beyond genomics to other domains requiring structured access to authoritative knowledge bases. More broadly, our results establish a principle for practical biomedical AI: connecting modest-sized LLMs to well-engineered, domain-specific tool environments is often more effective and cost-efficient than relying solely on ever-larger general-purpose models. In the context of rare disease research, this approach transforms a mature but complex database into an interactive assistant that handles technical barriers automatically while preserving MARRVEL’s curated evidence as the authoritative backbone and maintaining the essential role of expert oversight. The path forward involves improving accuracy through targeted fine-tuning, enhancing efficiency through architectural innovations to reduce token overhead, and expanding scope through multi-modal integration. MARRVEL-MCP demonstrates that the bottleneck in genomic AI is not model intelligence alone but the design of reasoning environments that make domain expertise executable—enabling sophisticated variant interpretation to become accessible to smaller, cheaper, more widely deployable models without sacrificing the rigor, transparency, and expert judgment that biomedical applications demand.

## 5 Acknowledgments

This work was supported by the Cancer Prevention and Research Institute of Texas (CPRIT, RP240131), the Chan Zuckerberg Initiative (2023-332162), the National Institutes of Health (NIH, U54NS093793), the Eunice Kennedy Shriver National Institute of Child Health and Human Development of the NIH (P50HD103555), the Chao Endowment, the Huffington Foundation, and the Jan and Dan Duncan Neurological Research Institute at Texas Children’s Hospital.

## References

[1] Sue Richards et al. “Standards and guidelines for the interpretation of sequence variants: a joint consensus recommendation of the American College of Medical Genetics and Genomics and the Association for Molecular Pathology”. en. In: Genetics in Medicine 17.5 (May 2015), pp. 405–424. ISSN: 10983600. DOI: 10.1038/gim.2015.30. URL: https://linkinghub.elsevier.com/retrieve/pii/S1098360021030318 (visited on 11/25/2025).

[2] Pragati Kore et al. “Improved allele frequencies in gnomAD through local ancestry inference”. en. In: Nature Communications 16.1 (Oct. 2025), p. 8734. ISSN: 2041-1723. DOI: 10.1038/s41467-025-63340-2. URL: https://www.nature.com/articles/s41467-025-63340-2 (visited on 11/26/2025).

[3] Julia Wang et al. “MARRVEL: Integration of Human and Model Organism Genetic Resources to Facilitate Functional Annotation of the Human Genome”. In: The American Journal of Human Genetics 100.6 (June 2017), pp. 843–853. ISSN: 0002-9297. DOI: 10.1016/j.ajhg.2017.04.010. URL: https://www.sciencedirect.com/science/article/pii/S0002929717301544 (visited on 11/25/2025).

[4] Melissa J. Landrum et al. “ClinVar: public archive of relationships among sequence variation and human phenotype”. en. In: Nucleic Acids Research 42.D1 (Jan. 2014), pp. D980–D985. ISSN: 0305-1048, 1362-4962. https://academic.oup.com/nar/article-lookup/doi/10.1093/ DOI:10.1093/nar/gkt1113nar/gkt1113. URL: https://www.sciencedirect.com/science/article/pii/S0002929717301544 (visited on 11/25/2025).

[5] Antonio Pedro Camargo et al. “Identification of mobile genetic elements with geNomad”. en. In: Nature Biotechnology 42.8 (Aug. 2024), pp. 1303–1312. ISSN: 1087-0156, 1546-1696. DOI: 10.1038/s41587-023-01953-y. URL: https://www.nature.com/articles/s41587-023-01953-y (visited on 11/25/2025).

[6] Joanna S Amberger et al. “OMIM.org: leveraging knowledge across phenotype–gene relationships”. en. In: Nucleic Acids Research 47.D1 (Jan. 2019), pp. D1038–D1043. ISSN: 0305-1048, 1362-4962. DOI: 10.1093/nar/gky1151. URL: https://academic.oup.com/nar/article/47/D1/D1038/5184722 (visited on 11/25/2025).

[7] Yanhui Hu et al. “An integrative approach to ortholog prediction for disease-focused and other functional studies”. en. In: BMC Bioinformatics 12.1 (Dec. 2011), p. 357. ISSN: 1471-2105. DOI: 10.1186/1471-2105-12-357. URL: https://bmcbioinformatics.biomedcentral.com/articles/10.1186/1471-2105-12-357 (visited on 11/25/2025).

[8] Damian Szklarczyk et al. “The STRING database in 2023: protein-protein association networks and functional enrichment analyses for any sequenced genome of interest”. eng. In: Nucleic Acids Research 51.D1 (Jan. 2023), pp. D638–D646. ISSN: 1362-4962. DOI: 10.1093/nar/gkac1000.

[9] Xiaoming Liu et al. “dbNSFP v4: a comprehensive database of transcript-specific functional predictions and annotations for human nonsynonymous and splice-site SNVs”. en. In: Genome Medicine 12.1 (Dec. 2020), p. 103. ISSN: 1756-994X. DOI: 10.1186/s13073-020-00803-9 URL: https://genomemedicine.biomedcentral.com/articles/10.1186/s13073-020-00803-9 (visited on 11/25/2025).

[10] Jeffrey R. MacDonald et al. “The Database of Genomic Variants: a curated collection of structural variation in the human genome”. en. In: Nucleic Acids Research 42.D1 (Jan. 2014), pp. D986–D992. ISSN: 0305-1048, 1362-4962. DOI: 10.1093/nar/gkt958. URL: https://academic.oup.com/nar/articlex-lookup/doi/10.1093/nar/gkt958 (visited on 11/25/2025).

[11] Eliete Da S. Rodrigues et al. “Variant-level matching for diagnosis and discovery: Challenges and opportunities”. en. In: Human Mutation (Mar. 2022), |phumu.24359. ISSN: 1059-7794, 1098-1004. DOI: 10.1002/humu.24359. URL: https://onlinelibrary.wiley.com/doi/10.1002/humu.24359 (visited on 11/25/2025).

[12] Michael A Gargano et al. “The Human Phenotype Ontology in 2024: phenotypes around the world”. en. In: Nucleic Acids Research 52.D1 (Jan. 2024), pp. D1333–D1346. ISSN: 0305-1048, 1362-4962. DOI: 10.1093/nar/gkad1005. URL: https://academic.oup.com/nar/article/52/D1/D1333/7416384 (visited on 11/25/2025).

[13] The GTEx Consortium et al. “The GTEx Consortium atlas of genetic regulatory effects across human tissues”. en. In: Science 369.6509 (Sept. 2020), pp. 1318–1330. ISSN: 0036-8075, 1095-9203. DOI: 10.1126/science.aaz1776. URL: https://www.science.org/doi/10.1126/science.aaz1776 (visited on 11/25/2025).

[14] Arzu Öztürk-Çolak et al. “FlyBase: updates to the Drosophila genes and genomes database”. en. In: GENETICS 227.1 (May 2024). Ed. by V Wood, iyad211. ISSN: 1943-2631. DOI: 10.1093/genetics/iyad211. URL: https://academic.oup.com/genetics/article/doi/10.1093/genetics/iyad211/7596147 (visited on 11/25/2025).

[15] Paul W Sternberg et al. “WormBase 2024: status and transitioning to Alliance infrastructure”. en. In: GENETICS 227.1 (May 2024). Ed. by V Wood, iyae050. ISSN: 1943-2631. DOI: 10.1093/genetics/iyae050. URL: https://academic.oup.com/genetics/article/doi/10.1093/genetics/iyae050/7640793 (visited on 11/25/2025).

[16] Richard M Baldarelli et al. “Mouse Genome Informatics: an integrated knowledgebase system for the laboratory mouse”. en. In: GENETICS 227.1 (May 2024). Ed. by V Wood, iyae031. ISSN: 1943-2631. DOI: 10.1093/genetics/iyae031. URL: https://academic.oup.com/genetics/article/doi/10.1093/genetics/iyae031/7635261 (visited on 11/25/2025).

[17] Eric W Sayers et al. “Database resources of the National Center for Biotechnology Information in 2025”. en. In: Nucleic Acids Research 53.D1 (Jan. 2025), pp. D20–D29. ISSN: 0305-1048, 1362-4962. DOI: 10.1093/nar/gkae979. URL: https://academic.oup.com/nar/article/53/D1/D20/7889242 (visited on 11/26/2025).

[18] Mohammed Toufiq et al. “Harnessing large language models (LLMs) for candidate gene prioritization and selection”. en. In: Journal of Translational Medicine 21.1 (Oct. 2023), p. 728. ISSN: 1479-5876. DOI: 10.1186/s12967-023-04576-8. URL: https://translational-medicine.biomedcentral.com/articles/10. 1186/s12967-023-04576-8 (visited on 11/26/2025).

[19] Model Context Protocol. URL: https://modelcontextprotocol.io/.

[20] Sean Kim and Raja Mazumder. Enhancing Scientific Reproducibility Through Automated BioCompute Object Creation Using Retrieval-Augmented Generation from Publications. Version Number: 1. 2024. DOI: 10.48550/ARXIV.2409.15076. URL: https://arxiv.org/abs/2409.15076 (visited on 11/25/2025).

[21] Manlian Bi et al. “BioRAGent: natural language biomedical querying with retrieval-augmented multiagent systems”. en. In: Briefings in Bioinformatics 26.5 (Aug. 2025), bbaf539. ISSN: 1467-5463, 1477-4054. DOI: 10.1093/bib/bbaf539. URL: https://academic.oup.com/bib/article/doi/10.1093/bib/bbaf539/8284869 (visited on 11/25/2025).

[22] Jiawei He et al. Retrieval-Augmented Generation in Biomedicine: A Survey of Technologies, Datasets, and Clinical Applications. Version Number: 3. 2025. DOI: 10.48550/ARXIV.2505.01146. URL: https://arxiv.org/abs/2505.01146 (visited on 11/25/2025).

[23] Xiangru Tang et al. MedAgents: Large Language Models as Collaborators for Zero-shot Medical Reasoning. 2311.10537 [cs]. June 2024. DOI: 10.48550/arXiv.2311.10537. URL: http://arxiv.org/abs/2311.10537 (visited on 11/25/2025).

[24] Nikita Mehandru et al. “BioAgents: Bridging the gap in bioinformatics analysis with multi-agent systems”. en. In: Scientific Reports 15.1 (Nov. 2025). Publisher: Nature Publishing Group, p. 39036. ISSN: 2045-2322. DOI: 10.1038/s41598-025-25919-z. URL: https://www.nature.com/articles/s41598-025-25919-z (visited on 11/25/2025).

[25] Hui Yang, Sifu Yue, and Yunzhong He. Auto-GPT for Online Decision Making: Benchmarks and Additional Opinions. 2306.02224 [cs]. June 2023. DOI: 10.48550/arXiv.2306.02224. URL: http://arxiv.org/abs/2306.02224 (visited on 11/25/2025).

[26] Shunyu Yao et al. ReAct: Synergizing Reasoning and Acting in Language Models. 2210.03629 [cs]. Mar. 2023. DOI: 10.48550/arXiv.2210.03629. URL: http://arxiv.org/abs/2210.03629 (visited on 11/25/2025).

[27] Lon Phan et al. “The evolution of dbSNP: 25 years of impact in genomic research”. eng. In: Nucleic Acids Research 53.D1 (Jan. 2025), pp. D925–D931. ISSN: 1362-4962. DOI: 10.1093/nar/gkae977.

[28] Jun Cheng et al. “Accurate proteome-wide missense variant effect prediction with AlphaMissense”. In: Science 381.6664 (Sept. 2023). ISSN: 1095-9203. DOI: 10.1126/science.adg7492. URL: http://dx.doi.org/10.1126/science.adg7492.

[29] Philipp Rentzsch et al. “CADD: predicting the deleteriousness of variants throughout the human genome”. In: Nucleic Acids Research 47.D1 (Oct. 2018), pp. D886–D894. ISSN: 1362-4962. DOI: 10.1093/nar/gky1016. URL: http://dx.doi.org/10.1093/nar/gky1016.

[30] Karthik A Jagadeesh et al. “M-CAP eliminates a majority of variants of uncertain significance in clinical exomes at high sensitivity”. In: Nature Genetics 48.12 (Oct. 2016), pp. 1581–1586. ISSN: 1546-1718. DOI: 10.1038/ng.3703. URL: http://dx.doi.org/10.1038/ng.3703.

[31] Keith J Kelleher et al. “Pharos 2023: an integrated resource for the understudied human proteome”. In: Nucleic Acids Research 51.D1 (Nov. 2022), pp. D1405–D1416. ISSN: 1362-4962. DOI: 10.1093/nar/gkac1033. URL: http://dx.doi.org/10.1093/nar/gkac1033

[32] Python Software Foundation. Python 3.13 Documentation. https://docs.python.org/3.13/. Accessed: 2025-11-26. 2025.

[33] FastMCP. FastMCP: A Fast, Pythonic Framework for MCP Tools. https://gofastmcp.com/. Accessed: 2025-11-26. 2025.

[34] GraphQL Foundation. GraphQL Specification. https://spec.graphql.org/October2021/. Accessed: 2025-11-26. 2021.

[35] Roy T. Fielding. “Architectural Styles and the Design of Network-based Software Architectures”. Defines the REST architectural style. PhD thesis. University of California, Irvine, 2000.

[36] LangChain AI. LangChain: A Framework for Building LLM Applications. https://github.com/langchain-ai/langchain. Accessed: 2025-11-26. 2025.

[37] Publiplots Developers. Publiplots: Publication-ready Plotting Library. https://pypi.org/project/publiplots/. Accessed: 2025-11-26. 2025.

[38] OpenRouter. OpenRouter: Unified API for AI Models. https://openrouter.ai/. Accessed: 2025-11-26. 2025.

[39] GGML and Contributors. llama.cpp: LLM Inference in C/C++. https://github.com/ggml-org/llama.cpp. Accessed: 2025-11-26. 2025.

[40] Angela R. Sung, Paolo Moretti, and Aziz Shaibani. “Case of late-onset Sandhoff disease due to a novel mutation in the HEXB gene”. en. In: Neurology Genetics 4.4 (Aug. 2018), e260. ISSN: 2376-7839. DOI: 10.1212/NXG.0000000000000260. URL: https://www.neurology.org/doi/10.1212/NXG.0000000000000260 (visited on 11/26/2025).

[41] Eric J. Topol. “High-performance medicine: the convergence of human and artificial intelligence”. en. In: Nature Medicine 25.1 (Jan. 2019). Publisher: Nature Publishing Group, pp. 44–56. ISSN: 1546-170X. DOI: 10.1038/s41591-018-0300-7. URL: https://www.nature.com/articles/s41591-018-0300-7 (visited on 11/25/2025).

