## Supplementary Materials for "MARRVEL-MCP: A Context-Engineered Natural-Language Query-to-Response Interface for Mendelian Disease Discovery"

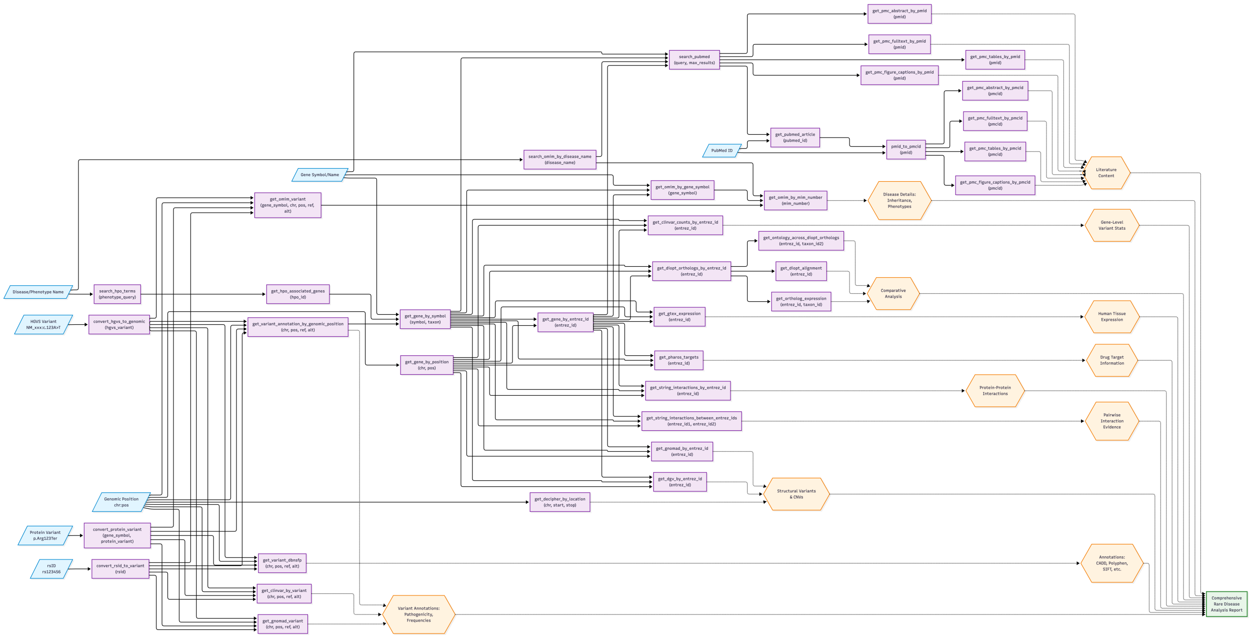


Figure S1: **Function-level view of the MARRVEL-MCP tool-chain, providing a detailed expansion of Figure 1.** This diagram maps the explicit API functions and data flows triggered from diverse user inputs (gene symbols, genomic positions, disease/phenotype terms, HGVS/rsID/protein variants, and literature identifiers) through granular modules, including variant conversion, gene information retrieval, ontology and ortholog analysis, expression profiling, interaction networks, disease annotation, and literature extraction (PubMed/PMC). Each functional node represents a specific callable operation, collectively illustrating how low-level functions are orchestrated to generate structured outputs that support comprehensive rare disease interpretation and evidence synthesis.

```python
@mcp.tool(
    name="get_clinvar_counts_by_entrez_id",
    description="Count all Clinvar variants for a given entrez gene id," + \
        "with subcounts for benign, likely benign, likely pathogenic, and pathogenic ClinVar designations.",
    meta={"category": "variant", "database": "ClinVar", "version": "1.0"},
)
async def get_clinvar_counts_by_entrez_id(entrez_id: str) -> str:
    try:
        data = await fetch_marrvel_data(
            f"""
            query MyQuery {{
                clinvarCountsByEntrezId(entrezId: {entrez_id}) {{
                    benign
                    likelyBenign
                    likelyPathogenic
                    pathogenic
                }}
            }}
            """
        )
        all_entries = await fetch_marrvel_data(
            f"""
            query MyQuery {{
                clinvarByGeneEntrezId(entrezId: {entrez_id}) {{
                    uid
                }}
            }}
            """
        )
        data_obj = json.loads(data)
        all_entries_obj = json.loads(all_entries)
        data_obj["data"]["clinvarCountsByEntrezId"]["all"] = len(
            all_entries_obj["data"]["clinvarByGeneEntrezId"]
        )
        data = json.dumps(data_obj, indent=2)
        return data
    except Exception as e:
        return json.dumps({"error": f"Failed to fetch data: {str(e)}"})
```

Figure S2: A Python code example demonstrating an example GraphQL-based query workflow for retrieving ClinVar variant counts by Entrez gene ID. The code illustrates the structural pattern of function calls and data flow without emphasizing specific MCP decorators, configuration details, or execution specifications.

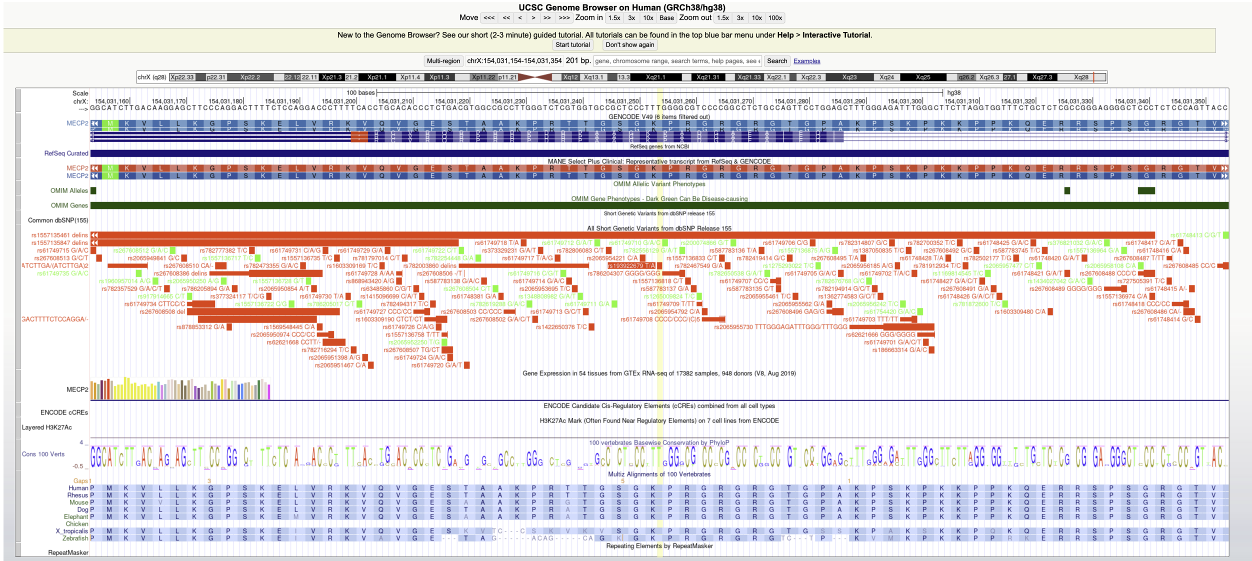


Figure S3: Genome browser view from MARRVEL illustrating the genomic context of rs193922679, a variant located within the MECP2 gene region. The visualization integrates multiple annotation tracks, including transcript structures, exon–intron boundaries, coding sequences, predicted variant consequences, and conservation scores. Red and green markers denote variant-level annotations and functional predictions derived from external databases, providing a consolidated view of potential biological impact across annotated isoforms. This figure highlights the structural and functional complexity of the MECP2 locus and underscores the importance of integrated visual representations for accurate interpretation of clinically relevant variants.

**Prompt**

For the gene HEXB, assess whether the variants c.298delC and G473S, found in a compound-heterozygous state, are likely to be pathogenic and sufficient to explain a Sandhoff-disease-like presentation. Provide a concise conclusion.

**Response**

The variant c.298delC (genomic chr5:74685557 delC) has no ClinVar or dbNSFP data available. This variant is a frameshift deletion, which is generally considered pathogenic. The variant G473S (genomic chr5:74718971 G>A) has conflicting interpretations of pathogenicity in ClinVar, with some submissions classifying it as "Likely pathogenic" and others as "Uncertain significance". The overall ClinVar significance is "Pathogenic/Likely pathogenic" with "criteria provided, multiple submitters, no conflicts". The dbNSFP scores for G473S show high pathogenicity predictions from CADD (Phred 36), REVEL (0.91498), and SIFT (deleterious). Given that c.298delC is a frameshift deletion, it is highly likely to be pathogenic. The G473S variant also has strong evidence for pathogenicity from ClinVar and dbNSFP. Therefore, the compound-heterozygous state of these two variants in HEXB is highly likely to be pathogenic and sufficient to explain a Sandhoff-disease-like presentation.

**Tool Calls**

get_gene_by_symbol, convert_protein_variant, convert_hgvs_to_genomic (x2), get_clinvar_by_variant (x2), get_variant_dbnsfp (x2)

Figure S4: The prompt, response, and tool calls for the HEXB variants interpretation.
